# Loss of Adaptive Capacity Drives Climate Vulnerability Across Taxonomic Scales in an Alpine Specialist Species Complex

**DOI:** 10.64898/2026.05.13.724772

**Authors:** Kristen Ruegg, Christen M. Bossu, Reza Goljani Amirkhiz, Nikunj Goel, Erica Robertson, Timothy M. Brown, Kathryn Bernier, Ben J. Vernasco, Peri E. Bolton, Erik R. Funk, Scott A. Taylor, Mevin B. Hooten, Erika S. Zavaleta

## Abstract

Accelerated warming at high elevations is having a disproportionate impact on alpine species. While assessments of climate vulnerability require quantifying the ecological and evolutionary components of adaptive capacity, such assessments are rare, especially in alpine systems. We leverage recent advances in population and landscape genomics to assess how variation in spatial heterogeneity and population connectivity across alpine systems influences adaptive capacity, using the North American Rosy-Finch species complex as a model system. In doing so, we clarify taxonomic relationships across the complex and identify one new ESU, the Sierra Nevada Rosy-Finch, based on its combined ecological and evolutionary distinctiveness. We then illustrate how combining genomic analyses with ecological data can improve estimates of adaptive capacity, sensitivity, and exposure and ultimately clarify climate vulnerability. Overall, our integrative analyses revealed that more isolated lineages, such as the Sierra Nevada Rosy-Finch, have lower adaptive capacity and face disproportionately high risks from climate change. This work highlights how conservation strategies that account for the multidimensional aspects of adaptive capacity can improve estimates of climate vulnerability.

## INTRODUCTION

Understanding the factors that determine species persistence amid rapid environmental change is a central goal of evolution and conservation biology. Climate change is driving rapid evolution, range shifts, and local extinction across species, leading some to argue that human activities are now the world’s greatest evolutionary force (Bumpus, 1898; Campbell-Staton et al., 2017; Ghalambor et al., 2015; Grant & Grant, 1993; Palumbi, 2001; Sinervo et al., 2010; Wiens, 2016). Alpine species may be particularly vulnerable to climate change because they occupy high-elevation habitats that may be experiencing accelerated, elevation-dependent warming, limiting opportunities for ecological buffering (Ernakovich et al., 2014; Grabherr et al., 2010; Mountain Research Initiative EDW Working Group, 2015). Species persistence under climate change depends on their capacity to shift distributions, adapt genetically to new conditions, or tolerate change through phenotypic plasticity (Beever et al., 2017). The ability to adapt genetically depends on a species evolutionary potential, which, in turn, is measured by genetic diversity, patterns of local adaptation, and the degree of population connectivity (Forester et al., 2022). Although incorporating evolutionary potential into climate vulnerability assessments across species is critical for effective conservation planning, such assessments remain rare, especially in alpine systems.

Climate vulnerability assessments typically involve measurements of their three main components: exposure, sensitivity, and adaptive capacity (Dawson et al., 2011; Foden et al., 2019; Thurman et al., 2020). Of these three components, adaptive capacity has historically been the most difficult to define due to its inherent complexity and confusion over its definition (Thompson et al., 2015; Thurman et al., 2020). Broadly, adaptive capacity includes a species’ dispersal ability, phenotypic plasticity, and evolutionary potential, the latter of which refers to its ability to adapt genetically to new environmental conditions (Forester et al., 2022). Recent advances in landscape genomics have facilitated estimates of genome-wide genetic diversity, gene flow, and local adaptation across heterogeneous landscapes, making it easier to assess evolutionary potential across a wide range of taxa (Aitken et al., 2024; Bernatchez et al., 2024). Further, the same genome-wide datasets can be used to estimate genomic offset, a framework that theoretically predicts the magnitude of allele-frequency change required for populations to adapt to future climatic conditions (Fitzpatrick & Keller, 2015; Forester et al., 2023; Rodriguez et al., 2025; Ruegg et al., 2018; but see Rellstab et al., 2021). In high-alpine systems, steep environmental gradients and differences in population connectivity and size may lead to substantial variation in evolutionary potential between populations, suggesting that adaptive capacity—and thus climate vulnerability—may vary substantially within a single species.

The rosy-finch species complex provides an excellent system for examining how environmental heterogeneity, range size, population connectivity, and local adaptation interact to shape evolutionary potential across alpine systems, because it comprises multiple taxa that differ across these variables. Rosy-finches are the highest-altitude breeding bird in North America, with nesting sites located in craggy, austere habitats in the Brooks, Rocky, Cascade, and Sierra Nevada Mountain ranges, as well as parts of the Aleutian Islands (Johnson et al., 2020; MacDougall-Shackleton et al., 2000). A century ago, Joseph Grinnell (1917) described the concept of the ecological niche using the Rosy-Finch as an example of a species whose distribution was limited by temperature. Recent modeling efforts predict that all North American Rosy-Finch species will experience dramatic range contractions due to climate change over the next 5-65 years (Conrad, 2015; Langham et al., 2015). Such declines may, in part, be due to the species’ dependence on snowpack, which in the Western U.S. has declined by 15-30% since the mid-20th century (Mote et al., 2018; Siirila-Woodburn et al., 2021). However, despite early work on this system, the remoteness of their alpine breeding habitats has left key aspects of their biology understudied, including some taxonomic relationships within the species complex and patterns of adaptive capacity within and among lineages (Johnson et al., 2020; Rosenberg et al., 2016).

The current accepted taxonomy of North American rosy-finches recognizes three species: the Black Rosy-Finch (*Leucosticte atrata*), Brown-capped Rosy-Finch (*L. australis*) (with no subspecies), and the Gray-crowned Rosy-Finch (*L. tephrocotis*), with several subspecies including the interior (*L. tephrocotis tephrocotis*), Hepburn’s (*L. t. littoralis*), Sierra Nevada (*L. t. dawsoni*), Wallowa (*L. t. wallowa*) and insular (Aleutian, *L. t. griseonucha*; and Pribilof, *L. t. umbrina*) Gray-crowned Rosy-Finches (Chesser et al., 2021). Taxonomic designations between some lineages within the Rosy-Finch complex remain challenging, largely because early phylogenetic reconstructions based on mtDNA provided little support for lineages described as distinct on morphological grounds (Drovetski et al., 2009) or did not include representatives of all recognized subspecies of Gray-crowned Rosy-Finch (Funk et al., 2021). Currently, the IUCN Red List classifies both the Black and Brown-capped Rosy-Finches as Endangered due to climate change and severe weather, but the Gray-crowned Rosy-Finch is listed as Least Concern (IUCN, 2026a, 2026b). However, the current lack of clarity regarding taxonomic relationships within the Gray-crowned Rosy-Finch may limit the ability to assess its vulnerability status. We quantify evolutionary relationships within this iconic species complex using expanded sampling across all major lineages to identify Evolutionarily Significant Units (ESUs) that may warrant conservation prioritization. We define ESUs as lineages exhibiting high genetic and ecological distinctiveness (Funk et al., 2021) and treat them as a taxonomic unit because of their widespread recognition among conservation practitioners responsible for implementing the Endangered Species Act.

To assess how adaptive potential and climate vulnerability vary between ESUs within this iconic alpine species complex, we leverage recent advances in population and landscape genomics. We first quantify patterns of connectivity and gene flow using whole-genome sequencing and increased sampling from previously unsampled regions. Our sampling includes all known species and subspecies, with particular emphasis on the two southernmost, dispersal-restricted taxa, the Brown-capped Rosy-Finch and the Sierra Nevada form of the Gray-crowned Rosy-Finch, hereafter referred to as the Sierra Nevada Rosy-Finch. After identifying ESUs, we assess how spatial heterogeneity and population connectivity impact evolutionary potential, measured as genetic diversity and effective population size, and apply genomic offset models to identify populations most at risk of maladaptation under future climate scenarios. We then demonstrate how integrating genomically based estimates of adaptive potential with ecological work on exposure and sensitivity can be used to more fully assess climate vulnerability and extinction risk in the two southernmost lineages.

## METHODS

### Sampling and Sequencing

A dataset comprising 203 Rosy-Finches from 27 sites across the United States (Figure 1, Table S1) was compiled from previously analyzed Rosy-Finch sequence data and new samples from the Sierra Nevada and Wallowa Rosy-Finch lineages. In summary, we included 115 individuals from 12 Brown-capped Rosy-Finch populations previously studied by DeSaix et al. (2022), 6 individuals from 2 Black Rosy-Finch populations, and 22 individuals from 7 Gray-crowned Rosy-Finch populations previously studied by Funk et al. (2021). We combined this with 52 new samples collected from 6 additional populations of the Sierra Nevada Rosy-Finch, and 8 individuals from a single population of the Wallowa Rosy-Finch in Oregon. One hundred and ninety two of these 203 samples were also used in Goel et al. (2026).

**Figure 1.**
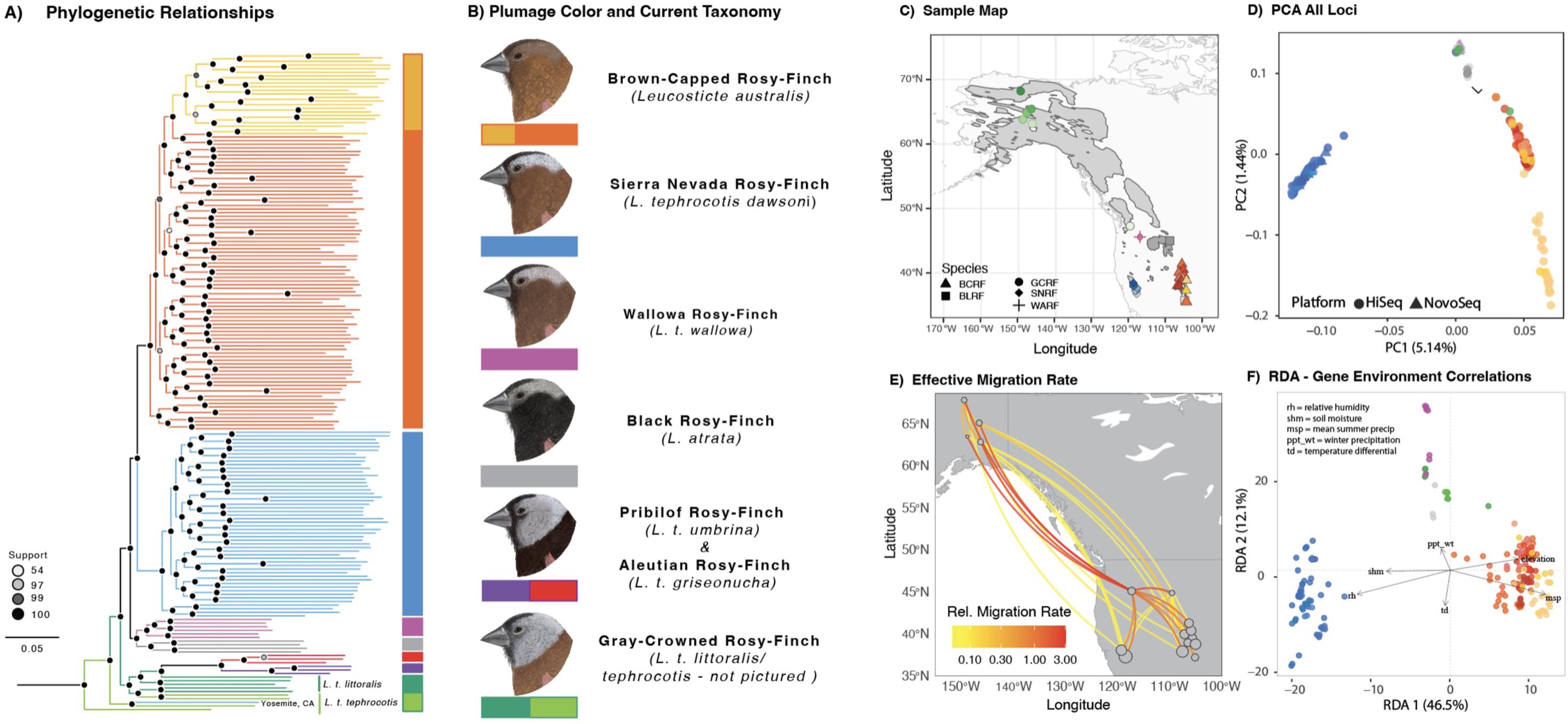
A) Phylogenetic Relationships within the Rosy Finch species complex. Current taxonomy suggests that the Sierra Nevada Rosy-Finch (Leucosticte tephrocotis dawsoni) is a subspecies of the Gray-Crowned Rosy-Finch (Leucosticte t. tephrocotis, light green and Leucosticte t. littoralis, dark green), but our results support it as its own Evolutionary Significant Unit. B) Plumage color variation and current taxonomic names within the Rosy Finch species complex. C) Relative migration rates between all sampling locations demonstrating ongoing gene flow across the complex, adapted from Goel et al (in press). D) PCA of the genetic relationships across all loci, showing that Sierra Nevada Rosy Finch is isolated along PC1, while the rest of the species in the complex follow an IBD pattern along PC2. E) Map of sampling locations in B and D. F) RDA showing how Sierra Nevada Rosy Finch has distinctly different gene environment correlations.

At all sites, Rosy-Finches were collected using potter traps and mist nets in June and July, when this species is most likely breeding (Theodosopoulos et al., 2023). Blood samples were taken from all birds via the brachial wing vein and stored in Queen’s lysis buffer at room temperature (Owen, 2011; Seutin et al., 1991). DNA was then extracted using the Qiagen DNeasy Blood and Tissue Kit with a modified protocol optimized to maximize DNA yield from feathers and quantified with the Qubit dsDNA HS Assay Kit (Thermo Fisher Scientific, USA). We used a modified version of Illumina’s Nextera Library Preparation protocol (Schweizer & DeSaix, 2023) to prepare whole-genome sequencing libraries and pooled the libraries by equal mass before sequencing. The pooled libraries were sequenced on two lanes of the Illumina NovaSeq 6000 at Novogene Corporation Inc.

### Bioinformatic Pipeline - Raw Reads to VCF

We provide an abbreviated version of the bioinformatics pipeline from raw reads to VCF creation and refer readers to Goel et al. (2026) for further details. Raw reads were processed using the mega-non-model-wgs-snakeflow pipeline and mapped to the Brown-capped Rosy-Finch genome assembly (*Leucosticte australis*, GCA_025504685.1). Reads were trimmed with fastp (Chen et al., 2018) and aligned using BWA-MEM2 (Li & Durbin, 2009; Vasimuddin et al., 2019). Alignments were processed with SAMtools (Danecek et al., 2021; Li et al., 2009) and Picard to sort BAM files, add read groups, and mark PCR duplicates. Sequencing depth was calculated with SAMtools, and BAM files were downsampled to 5× coverage to reduce depth variation (Lou & Therkildsen, 2022). Variants were called using HaplotypeCaller in GATK v4.2.6.1 (McKenna et al., 2010; Van der Auwera et al., 2013). Variants were filtered following GATK best practices to remove systematic errors (Van der Auwera et al., 2013): QD < 2.0 || FS > 60.0 || MQ < 40.0 || MQRankSum < -12.5 || ReadPosRankSum < -8.0. We further filtered on missingness and quality, retaining biallelic SNPs with < 20% missing data, MAF ≥ 0.05, and QUAL > 30. To minimize sequencing-platform effects (HiSeq vs. NovaSeq), we conducted an F_ST_ outlier scan with VCFtools (Danecek et al., 2011) with individuals from White Mountain sequenced on both platforms and removed loci with F_ST_ > 0.05. Missing loci were imputed with Beagle version 4.1, which uses the genotype likelihood directly from non-missing loci(Browning & Browning, 2016), and confirmed that a PCA generated with and without imputation did not alter population structure. We used plink (--indep-pairwise 50 5 0.2) to prune our whole-genome sequence data for linkage disequilibrium (Chang et al., 2015; Purcell et al., 2007). We generated a second dataset to estimate the effective population size that was not MAF-filtered but was filtered for minor allele count (MAC >=2), and missing genotypes were imputed using Beagle 4.1 (Browning & Browning, 2016). This dataset, hereafter called the MAC filtered dataset, retained rare alleles and linked loci and was used to estimate effective population size.

### Bioinformatic Pipeline - BAMS to Genotype Likelihoods

We used a second Snakemake workflow pipeline (Mölder et al., 2021) called loco-pipe (Zhou et al., 2024) to calculate genotype likelihoods in lieu of called genotypes. The downsampled BAM files from the mega-non-model-wgs-snakeflow pipeline were used as input into the loco-pipe workflow (Zhou et al., 2024). We generated a separate “unit” file for each sequencing platform (HiSeq vs. NovaSeq) that linked the BAM filename to the sample name and population, since we ran this workflow separately for samples sequenced on each platform. Within the loco-pipe workflow, global depth was estimated, and global SNP calling for all breeding birds was completed using ANGSD to generate genotype likelihoods (Korneliussen et al., 2014). We implemented variable base quality mapping parameters for NovaSeq and HiSeq (minq: 30 and 20, respectively), however the same minimum mapping quality score was used (minmapq: 30). In both workflows, at least 50 percent of individuals were required to have at least one read per site (minind_proportion 0.5 and mindepthind 1), a minimum allele frequency threshold (maf: 0.05), a threshold for a site to be considered as a SNP (1e-6), and we removed triallelic variants. We also computed an extended per-base alignment quality (BAQ) to directly estimate the probability of misalignment around indels (baq: 2) (Li, 2011). To remove linked loci, we subsample the global SNP set, selecting 1 SNP out of every 10.

### ESU Delineation

To construct a Rosy-Finch phylogeny, we used samples from the Aleutian and Pribilof Islands, along with six Asian Rosy-Finch samples previously sequenced in Rosy-Finch phylogenomic analyses (Funk et al., 2021), as outgroup lineages. The mega-non-model-snakeflow pipeline was run from the beginning to call variants (including new variants from these new populations), applying filters for systematic errors, platform effects, quality, and missingness. We used plink (--indep-pairwise 50 5 0.5) to prune our whole-genome sequence data for linkage disequilibrium (Chang et al., 2015; Purcell et al., 2007). We also added a more stringent individual filter by identifying individuals with more than 20% missing called genotypes using VCFtools *–missing* (Danecek et al., 2011) and removing them from the VCF file using BCFtools (Danecek et al., 2021). We then removed an admixed Brown-capped Rosy-Finch individual. We converted the individual and LD-pruned VCF to a phylip file using a custom Python script, vcf2phylip.py (Ortiz, 2019), randomly resolving heterozygous sites to avoid IUPAC ambiguities. Then, we constructed the maximum-likelihood phylogeny in IQ-TREE (Minh et al., 2020) using the GTR+F+ASC model. The analysis was performed using the ultrafast bootstrap approximation (-B 1000; Hoang et al., 2018) to reduce the risk of overestimating branch support, with hill-climbing nearest-neighbor interchange (-bnni). Finally, the tree was visualized in FigTree 1.4.4 (Rambaut, 2018).

To determine the number of genetically distinct breeding clusters within the continental Rosy-Finch complex, we estimated individual admixture proportions by running five replicate runs of PCA_NGSD_ –admix (Meisner & Albrechtsen, 2018) on the LD pruned imputed dataset for *K* values 2-7, manually selecting the number of eigenvalues (-e) for modelling of individual allele frequencies and identified consistent geographic signatures of clustering across highly supported runs. We performed three hierarchical admixture analyses: (1) all continental Rosy-Finch lineages, (2) only populations that clustered with the Gray-Crowned Rosy-Finch in the first run, then (3) only the Brown-Capped Rosy-Finches. Using R 4.5.2 (R Core Team 2025), we visualized breeding clusters on a base map from Natural Earth (https://naturalearthdata.com), clipped to the breeding ranges of each species, and scaled the color transparency according to the posterior probability of group membership from the individual admixture proportions (Figure 2). We used eBird shapefiles for the breeding ranges of Black Rosy-Finch and Gray-crowned Rosy-Finch, and, given the limited distribution of the Wallowa Rosy-Finch, we created a polygon buffer around the specific sampling location to facilitate color deposition. For the Sierra Nevada Rosy-Finch and Brown-capped Rosy-Finch, we used the predicted ranges based on habitat suitability (Amirkhiz et al., 2025, 2026), excluding unsuitable habitats (see Climate heterogeneity in BCRF and SNRF below).

**Figure 2.**
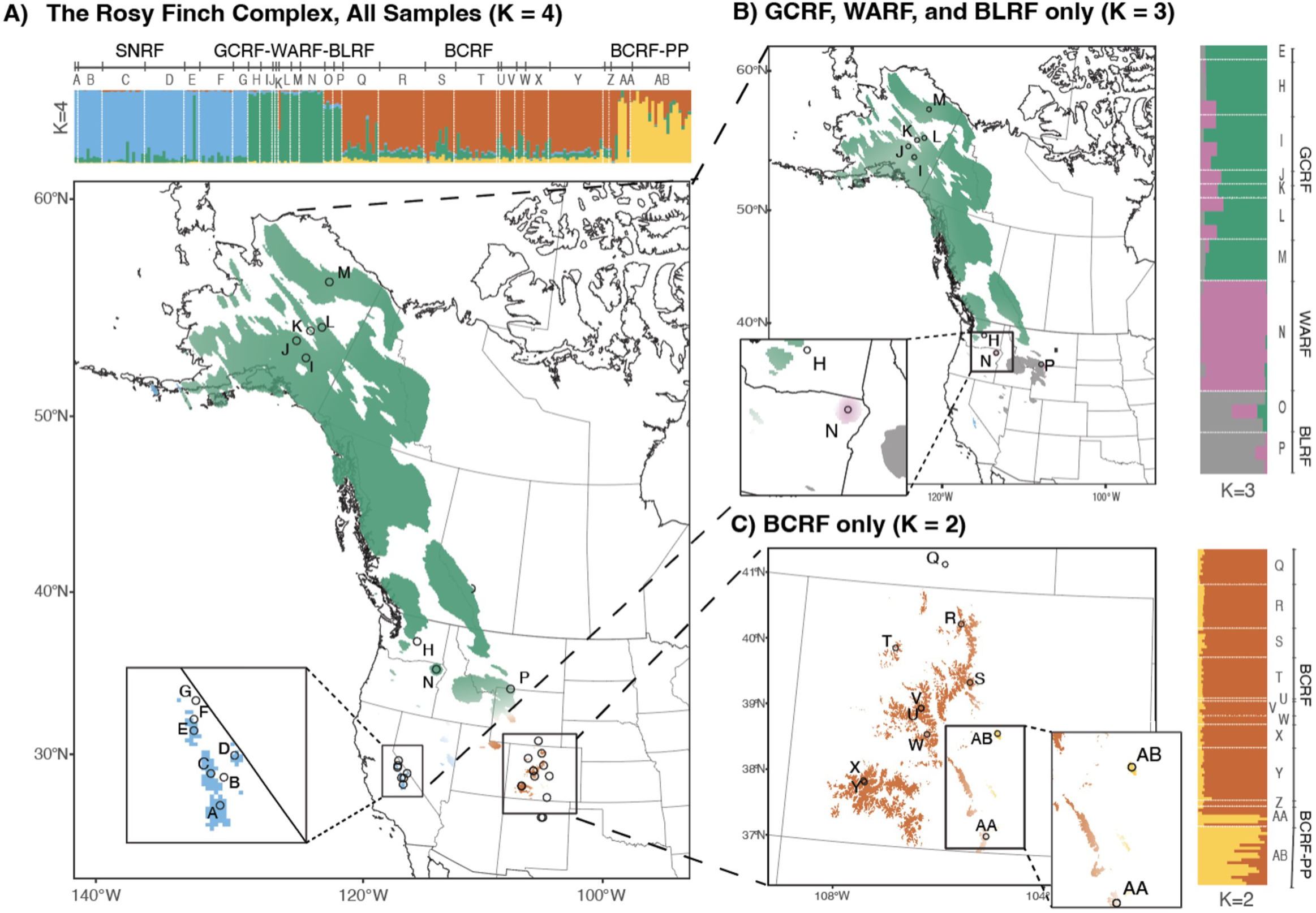
Hierarchical admixture plot generated using PCAngsd –admix. A) All samples within the species complex, showing that the first 4 groups that come out are the Sierra Nevada Rosy-Finch (SNRF), the Gray-crowned Rosy-Finch (GCRF), and the Brown-capped Rosy-Finch (BCRF), with the Pikes Peak (BCRF_PP) location coming out as separate. B) Admixture without SNRF and BCRF revealed additional structure within the GCRF, including Wallowa (WARF), and the Black Rosy Finch (BLRF), which is currently designated as a separate species. C) BCRF only, showing the location of the Pikes Peak population in the southeast corner of the range.

### Evolutionary Potential

As part of the loco-pipe workflow, we also estimated inbreeding (F_IS_) and individual-level heterozygosity (H_IND_). To quantify inbreeding, or deviations from HWE as an excess or deficiency in observed heterozygotes, we used an inbreeding coefficient statistic implemented in PCA_NGSD_ (Meisner & Albrechtsen, 2019). The method incorporates individual allele frequencies to account for population structure. We also calculated individual-level observed heterozygosity using ANGSD --het, estimating the site frequency spectrum per individual (excluding the sex chromosomes) and the proportion of heterozygous genotypes. We checked and corrected for putative sequencing-platform effects on heterozygosity using a linear mixed model in the R package lme4 (Bates et al., 2015). We modeled individual heterozygosity values given random site and sample effects and a fixed sequencing platform effect (het ∼ Platform + (1 | sample_name) + (1 | Population)). We added 84 newly sequenced Brown-capped Rosy-Finch individuals on a single NovaSeq lane (*i.e.,* duplicates of individuals sequenced on HiSeq lanes in the current analyses) to the 19 White Mountain, CA individuals sequenced on both platforms in this analysis (including 4 duplicate samples) to strengthen the regression and calculate the coefficient associated with sequencing platform and correct the heterozygosity estimates. We additionally assessed and corrected heterozygosity at candidate regions, defined as 500 bp regions surrounding 683 candidate loci identified using a hierarchical Bayesian GEA model of Rosy-Finches that accounted for migration (Goel et al., 2026). We plotted spatial interpolation of the global and candidate H_IND_ using custom R scripts, excluding the non-breeding populations of Fairbanks, Ellensburg, and Elk Mountain.

### Estimating N_e_ using GONE

We estimated the recent demographic history of Brown-capped and Sierra Nevada Rosy-Finches using the program GONE, which effectively accounts for variable sequencing depth across the genome, allowing for differences in sequencing depth both between individuals and within genomes (Santiago et al., 2020). This method leverages the fact that observed linkage disequilibrium (LD) at different genetic distances provides information about past *N*_e_ at different generations. To estimate the effective population sizes (N_e_) of Brown-capped Rosy-Finch and Sierra Nevada Rosy-Finch, we used the MAC-filtered imputed dataset and filtered out individuals with more than 17% missing called genotypes. Because GONE assumes panmixia, we subset the data into three lineage-specific datasets: 1) Sierra Nevada Rosy-Finch, 2) the main Brown-capped Rosy-Finch cluster, and 3) the smaller Brown-capped cluster that included individuals from Pikes Peak and several individuals from Mountain Maxwell. We subsequently removed variants not in Hardy-Weinberg equilibrium using BCFtools (Danecek et al., 2021) and removed chromosomes with fewer than 50,000 SNPs and/or less than 20cM. We performed 100 replicate analyses of GONE (Santiago et al., 2020), randomly sampling 50,000 SNPs per autosome, over 150 generations. We plotted the combined GONE results in R (R Core Team 2025) using custom scripts to visualize the median N_e_ across replicates and the 95% confidence intervals. We used the average generation time for Gray-crowned (gt=2.19) and Brown-capped Rosy-Finch (gt=2.04; Bird et al., 2020), and the average year of sampling collection in this study, 2019, to plot the change in effective population time in years.

### Habitat Suitability and Heterogeneity

To compare climatic heterogeneity between Brown-capped Rosy-Finch and Sierra Nevada Rosy-Finch study areas, we used two complementary summaries of dispersion in a shared environmental space. First, we calculated descriptive dispersion metrics in the PCA space, including the variance along PC1 and PC2, the total variance across the first two axes, and the area of the 95% covariance ellipse. These metrics summarize how broadly each study area occupies the ordination space, and ellipse-based summaries are a standard geometric representation of multivariate spread (Friendly et al., 2013). Because the PCA was built from a shared set of bioclimatic variables extracted from suitable-habitat cells, these summaries describe relative climatic breadth in a common environmental space (Broennimann et al., 2012). Second, we used a permutation-based centroid-distance analysis to formally compare climatic heterogeneity between study areas. Suitable-habitat cells were represented in standardized multivariate climate space, and climatic heterogeneity was quantified as the mean Euclidean distance of cells to the centroid of their study area. A greater mean distance to the centroid indicates greater within-group multivariate dispersion, consistent with the framework of Anderson (2006) and Anderson et al. (2006). We then compared the observed difference in mean distance to the centroid between study areas against a null distribution generated by permuting study-area labels.

### Adaptive Index & Genomic Offset

Previous Rosy-Finch genotype-environment association analyses (Goel et al., 2026) identified 6 least correlated environmental variables important to the genetic structure of Rosy-Finches: (a) Soil water content (SWC), (b) mean summer precipitation (MSP), (c) relative humidity (RH), (d) summer heat moisture index (SHM), (e) continentality (TD, or seasonality), and (f) mean July snowpack. However, mean July snowpack and soil water content do not have future climate change projections, so we substituted correlated AdaptWest (AdaptWest Project, 2022) bioclimate variables (*e.g.,* winter precipitation (PPT_wnt) and elevation). We performed redundancy analyses (RDA) using these variables to calculate adaptive indices and genomic offset. An adaptive index ultimately defines how adaptive genetic variation is distributed across landscapes by quantifying how genetic markers correlate with bioclimate variables and how those variables correlate with one another (Capblancq & Forester, 2021). We calculated genomic offset to illustrate a potential shift in the adaptive optimum of these top variables induced by climate change (Capblancq et al., 2020). We estimated genomic offsets for Rosy-Finches by applying the same procedure used for the current climate to two future climate projections, SSP2-45 and SSP5-85 (Wang et al., 2016), which differ in the trajectory of greenhouse gas emissions and thus projected climatic changes for 2041-2070, and predicted adaptive indices based on RDA1 and RDA2. We calculated the Euclidean distance between the current and future adaptive indices along the first two RDA axes, yielding a genomic offset value. Genomic offsets estimated for each future scenario were mapped across the species’ range to identify areas where the greatest allelic change must occur under future climate change, highlighting putative high-risk areas.

## RESULTS

### ESU delineation

To identify ESU within the continental Rosy-Finch complex, we used a multi-faceted approach, leveraging phylogenomic inference, admixture estimates, principal component analysis, and redundancy analyses to assess how this alpine specialist varies genetically across environmental space. The average sequencing coverage of individuals before down-sampling was 6.58X (range: 1.5-23.12X). Overall, we identified 3.87 million high-quality variants after filtering for quality, missingness, and batch effects, and conducted phylogenomic analyses with IQ-Tree on 189 individuals and 2,826,075 unlinked variant sites. One Russian individual, 18N01242, was designated as the outgroup taxon; however, 5 additional Asian Rosy-Finch individuals and 6 Island Rosy-Finch subspecies (3 Aleutian and 3 Pribilof samples) were included in total. The consensus tree, constructed from 1000 bootstrap trees, showed strong support for a monophyletic North American clade (>95% support; Figure 1a). Our results recovered Gray-crowned Rosy-Finch as sister to the remaining Rosy-Finch lineages; although it was paraphyletic, and we could not resolve the relationship between *L. t. tephrocotis* and *L. t. littoralis* subspecies. All other main Rosy-Finch lineages were strongly supported (>95%; Figure 1a).

Contrary to previous analyses by Funk et al. (2021), our results supported the nesting of the Aleutian and Pribilof Gray-crowned Rosy-Finch subspecies within the Gray-crowned lineage, rather than sister to the mainland Gray-crowned lineages. However, like Funk et al. (2021), within the clade of island individuals, the two currently recognized subspecies from the Aleutian and Pribilof islands were recovered as reciprocally monophyletic. Our results further support that Wallowa and Black Rosy-Finches are reciprocally monophyletic (Figure 1a) and sister to the clade containing Brown-capped and Sierra Nevada Rosy-Finches, which are also reciprocally monophyletic. Nested within the monophyletic Brown-capped Rosy-Finch clade, Pikes Peak individuals and several individuals from Mountain Maxwell formed a paraphyletic clade. The one exception to the patterns described above is the single Yosemite individual, also used in Funk et al. (2021), sampled within the Sierra Nevada Rosy-Finch range, but clustering with the Gray-crowed Rosy-Finch clade. Given the absence of an admixture signal between this individual and other Sierra Nevada Rosy-Finches, we hypothesize that it is either a misidentified sample, a migrant, a Gray-crowned Rosy-Finch, or the result of a laboratory error.

The previously estimated migration by Goel et al. (2026) and a principal component analysis based on imputed genomic data from the continental lineages supported the phylogenomic analyses above. The PCA analysis revealed that the Sierra Nevada Rosy Finch was isolated from the rest of the complex along PC1, which explained 5.14% of the genetic variation, while the rest of the complex separated along PC2, which explained 1.44% of the variance (Figure 1d). These results were mirrored by the lower effective migration rates into the Sierra Nevada Rosy-Finch found in Goel et al. (2026) (Figure 1e). Redundancy analyses indicate that the Sierra Nevada Rosy-Finches separated from the rest of the complex along RDA1, which explained 46.5% of the variance and was linked to four climatic variables: mean summer precipitation, elevation, relative humidity, and the summer heat moisture index. In contrast, RDA2 separated the remaining lineages, explained 12.1% of the variance, and was linked to winter precipitation and continentality.

Hierarchical admixture analyses of all unlinked loci (adaptive and non-adaptive) and all continental individuals identified four main genetic clusters, the most differentiated genetic groups: Sierra Nevada Rosy-Finch, Gray-Crowned Rosy-Finch, Brown-capped Rosy-Finch, and Pikes Peak Brown-capped Rosy-Finch (Figure 2a, b, & c). It was not until the exclusion of the highly differentiated lineages that additional population structure became apparent, and we identified clear clustering of Gray-crowned, Wallowa, and Black Rosy-Finches (Figure 2b). However, this may be related to the low sample size for these lineages.

### Evolutionary Potential

To quantify evolutionary potential across the range, we used the loco-pipe workflow to estimate individual-level heterozygosity (H_IND_) from genotype likelihoods, thereby avoiding potential imputation-related artifacts. We plotted heterozygosity across the landscape and observed a decreasing trend from the southern Rockies (Brown-capped Rosy-Finch) toward the northernmost part of the breeding range (Gray-crowned Rosy-Finch in Alaska); however, Sierra Nevada Rosy-Finch populations exhibited the lowest heterozygosity estimates (Figure 3a). To further quantify evolutionary potential at adaptive loci and compare it to all loci, we localized H_IND_Adaptive_ estimates at candidate regions and found that they were relatively equal across the Rosy-Finch range, except for the Sierra Nevada Rosy-Finch populations, which had appreciably lower heterozygosity estimates than the other populations (Figure 3b).

**Figure 3.**
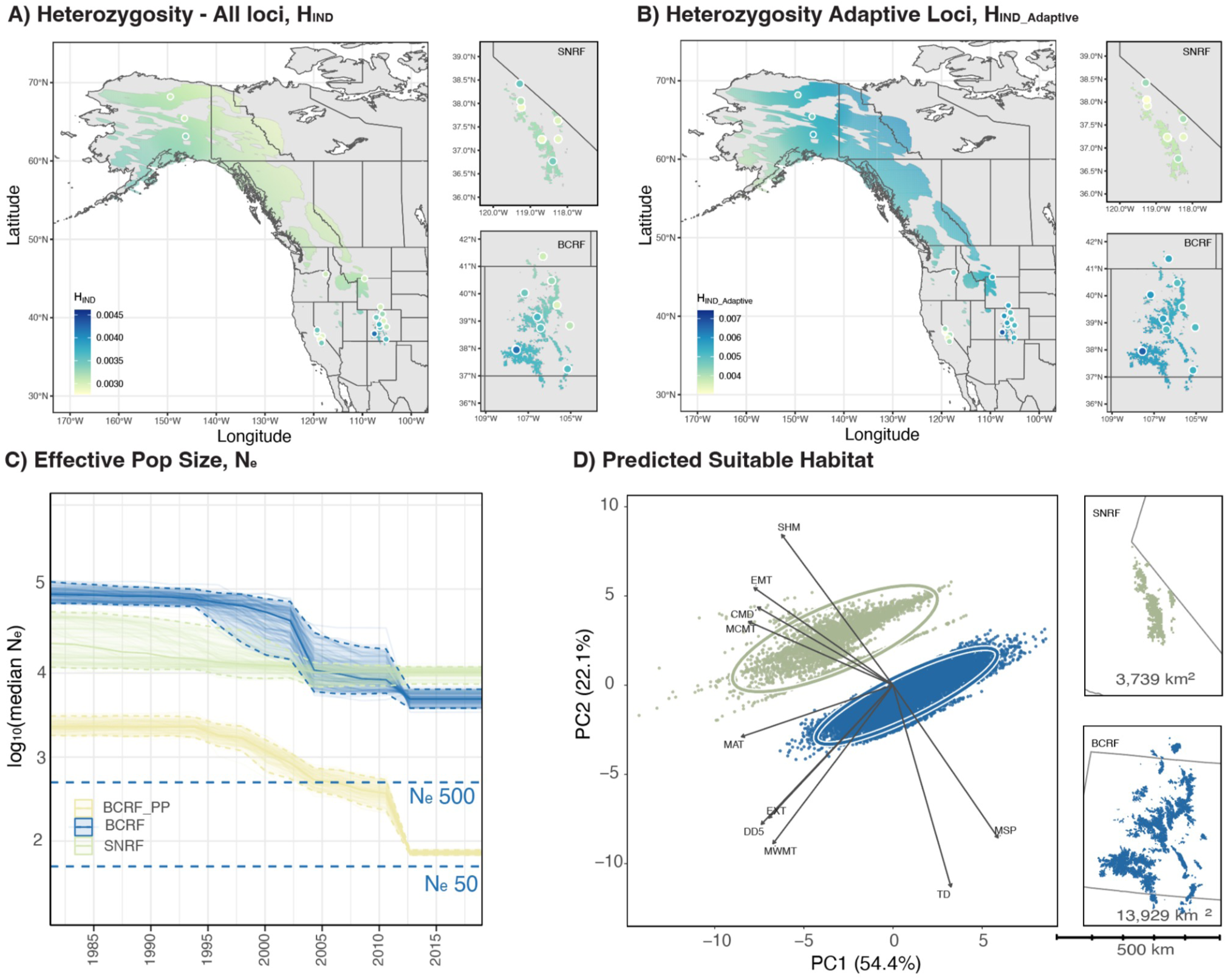
Evolutionary Potential. A) Individual heterozygosity shows lower heterozygosity in the south than in the north, with Sierra Nevada Rosy-Finch (SNRF) having lower heterozygosity than Brown-capped Rosy-Finch (BCRF). B) Individual heterozygosity at candidate loci identified by RDA demonstrates the lowest heterozygosity in the Sierra Nevada Rosy-Finch. C) Ne through time, estimated using the program GONE, indicates that all three populations have declined in recent decades, with SNRF and BCRF remaining above the threshold of 500 individuals recommended for maintaining evolutionary potential, while BCRF in Pikes Peak is nearing the threshold of 50 individuals, suggesting likely inbreeding. D) Principal component analysis (PCA) of bioclimatic conditions, extracted from predicted suitable habitat cells, shows points representing sampled suitable-habitat cells, ellipses indicating 95% covariance, and arrows illustrating loadings of the bioclimatic variables: TD = temperature difference, MSP = mean summer precipitation, MWMW = mean temperature of the warmest month, DD5 = degree-days above 5°C, EXT = extreme max temperature over 30 years, MAT = mean annual temperature, MCMT = mean temperature of the coldest month, CMD = climate moisture index, EMT = extreme min temperature over 30 years, SHM = summer heat moisture index. Maps display the predicted suitable habitat area for the SNRF and BCRF, shown at the same spatial scale.

### Effective Population Size using GONE

Estimates of effective population size, based on linkage equilibrium, generally supported a decline in the effective population sizes of both the main Brown-capped and Sierra Nevada Rosy-Finch approximately 20 years ago (Figure 3c), though the patterns of decline differ. The main Brown-capped Rosy-Finch population historically had a larger effective population size and experienced a steeper decline than the Sierra Nevada population. The Pikes Peak Brown-capped Rosy-Finch group began declining around the same time, with a marked decline around 2010. Estimates of effective population size at time of sampling show a lower median N_e_ estimate in the main Brown-capped Rosy-Finch (N_e_=4,923, 95% CI: 3,823 - 6,422) compared to Sierra Nevada Rosy-Finch estimates (N_e_=10,266, 95% CI: 7,411 - 11,855). Both estimates exceed the N_e_=500 proposed for maintaining evolutionary potential in the long term (Franklin, 1980) and the recommended doubling (N_e_=1000) based on a theoretical and empirical review of this conservation rule (Frankham et al., 2014). The Pikes Peak lineage of Brown-capped Rosy-Finch had the lowest effective population size estimate (N_e_=73, 95% CI: 68-78) and lies on the cusp of the N_e_=50 conservation rule, where extinction risk is high (Franklin, 1980).

### Habitat Suitability and Heterogeneity

Both the descriptive summaries and the permutation-based centroid-distance analysis indicated greater climatic heterogeneity in the Sierra Nevada Rosy-Finch range, spanning the Sierra Nevada, White, and Sweetwater Mountains, than in the Colorado Rockies. In the descriptive PCA-space summaries, SNRF had higher total variance than BCRF (7.25 vs. 5.18) and a substantially larger 95% ellipse area (37.22 vs. 18.65), indicating a broader spread along the first two PCA axes. The two study areas were also strongly separated in climate space, with a centroid distance of 5.21 and no convex-hull overlap (Jaccard = 0). These descriptive summaries, therefore, suggest that SNRF occupies a broader and more distinct climatic space in the plotted ordination. The formal centroid-distance analysis reached the same conclusion. Mean distance to centroid was higher for SNRF than for BCRF (2.97 vs. 2.53), corresponding to an observed difference of 0.44, or roughly 17% greater climatic dispersion in SNRF. None of the 9,999 permutations produced a difference as large as the observed one, indicating strong support for greater climatic heterogeneity in SNRF. This result means that suitable-habitat climates in the Sierra Nevada and adjacent ranges are more dispersed around their multivariate centroid than those in the Colorado Rockies.

### Adaptive Index & Genomic Offset

Initial attempts to visualize genomic offset and adaptive indices across the entire Rosy-Finch breeding range reveal highly negative adaptive values of RDA1 along coastal Alaska where breeding records are less certain relative to the rest of the range (Figure 4a). This bias likely arises because the eBird distribution map of the Gray-crowned Rosy-Finch is inaccurate (e.g., too broad) for an alpine specialist with a very limited range, leading the model to sample environmental predictors in areas that are not actually part of the species’ more restricted range. Therefore, to more accurately compare adaptive indices and genomic offsets across the ranges of the Brown-capped and Sierra Nevada Rosy-Finches, we used range maps generated from habitat-suitability models by Amirkhiz et al. (2025, 2026). After removing putative biases caused from using an overly broad range map, we demonstrated that adaptive indices clearly differ between Brown-capped Rosy-Finch in the Rockies (negative RDA1 scores) and Sierra Nevada Rosy-Finch in the Sierras (positive RDA1 scores). When these relationships are projected onto future climate-change models in the genomic offset portion of the analysis, we see that the Sierra Nevada Rosy-Finch range has higher genomic offset levels than the Brown-capped Rosy-Finch (Figure 4b) which is in keeping with its lower evolutionary potential.

**Figure 4.**
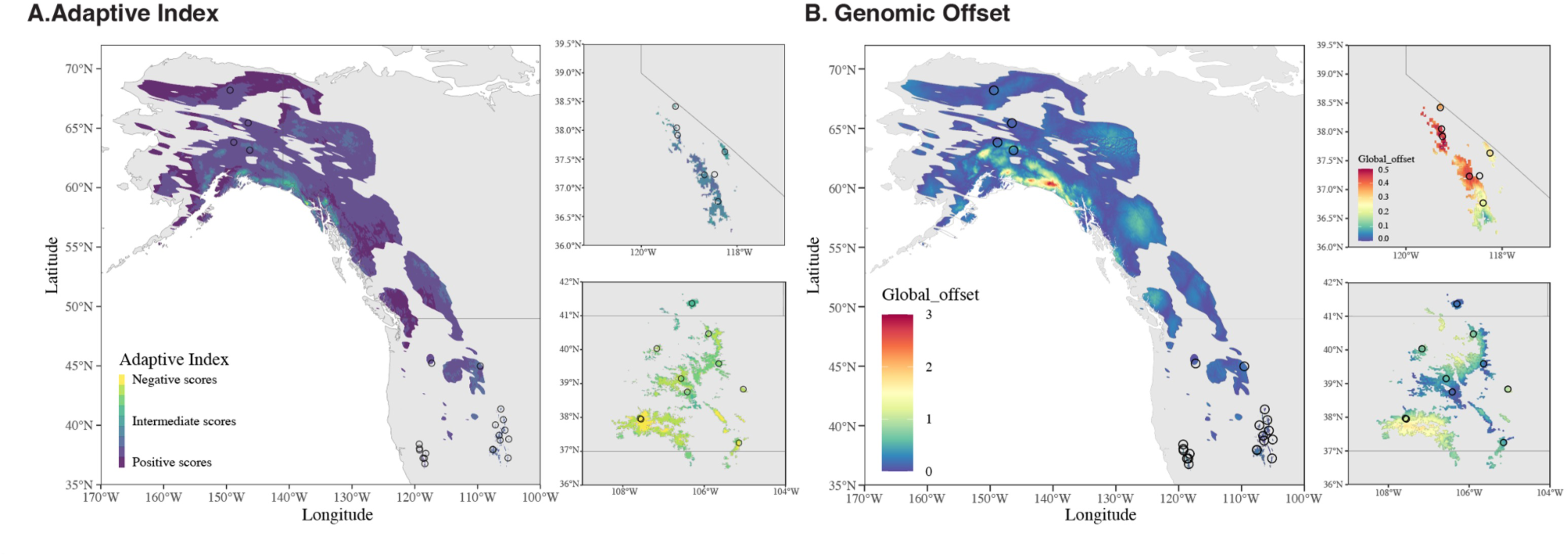
A) Adaptive Index from Redundancy Analysis with close-ups of the Brown-capped and Sierra Nevada Rosy-Finch ranges. Positive values in the Sierra Nevada Mountains and negative values in the Rockies support the idea that Sierra Nevada and Brown-capped Rosy-Finches are adapted to different climates. B) Genomic offset using a moderate climate change scenario (SSP2-45) supports the idea that Sierra Nevada Rosy Finches will have a harder time adapting to future climate change.

## DISCUSSION

Alpine species may be particularly susceptible to climate change because they inhabit high-elevation environments where warming is often accelerated, resulting in a reduction in suitable habitat and limited dispersal opportunities (Ernakovich et al., 2014; Grabherr et al., 2010; Mountain Research Initiative EDW Working Group, 2015). Measurements of adaptive capacity, including dispersal ability, phenotypic plasticity, and evolutionary potential, are central to assessing vulnerability to climate change but have historically been difficult to define (Exposito-Alonso et al., 2022; Thompson et al., 2015; Thurman et al., 2020). We use the Rosy-Finch species complex as a model to examine how variation in spatial heterogeneity and population connectivity across alpine systems may influence adaptive capacity. In doing so, we clarified taxonomic relationships across the Rosy-Finch species complex and identified one new ESU, the Sierra Nevada Rosy-Finch, based on its combined ecological and evolutionary distinctiveness. We then combined our genomic estimates of evolutionary potential with information on exposure and sensitivity to assess how climate vulnerability varies with adaptive capacity.

### Evolutionary Significant Units

Evolutionary relationships within the Rosy-Finch species complex have historically been difficult to resolve. While the original species classifications were largely based on plumage differences, Drovetski et al. (2009) detected no phylogenetic structure within the complex using mtDNA sequencing, leading them to propose treating the taxa as a single species. Later, Funk et al. (2021) used whole-genome sequencing to assess relationships across the complex and found phylogenetic support for Black and Brown-capped Rosy-Finches as distinct lineages, but Gray-crowned Rosy-Finches remained paraphyletic. We use whole-genome sequencing and increased sampling from previously understudied regions and find that our results are largely concordant with those of Funk et al. (2021), with a few key differences.

The first and most striking difference between our results and previous work is the genetic and ecological distinctiveness of the Sierra Nevada Rosy-Finch (*L. tephrocotis dawsonii*), which is currently considered a subspecies of the Gray-crowned Rosy-Finch. Whereas the placement of the Sierra Nevada lineage in Funk et al. (2021) was less clear, our analyses suggest that it is genetically and ecologically divergent from all other Rosy-Finches. Within our PCA, the Sierra Nevada Rosy-Finch is the most divergent lineage along PC1 (5.71% variance), while all other lineages within the complex follow an isolation-by-distance pattern along PC2 (1.73% variance) (Figure 1d). The increased isolation of the Sierra Nevada Rosy-Finch is also mirrored in the phylogenetic, effective migration rate, and admixture results (Figure 1a, e, & 2a). Further, habitat suitability and GEA analyses suggest that Sierra Nevada Rosy-Finches occupy and are adapted to the distinctive Mediterranean alpine climate of the Sierra Nevada Mountains, characterized by mild winters and limited summer rainfall relative to most other alpine regions across the world (Figure 1d, e, & f & Figure 4a; Rundel, 2011). Together, the ecological and genetic distinctiveness of the Sierra Nevada Rosy-Finch suggest it should be considered a distinct ESU for conservation purposes. These results are especially timely given recent work by Amirkhiz et al. (2025), showing that climate change has already reduced suitable habitat for the Sierra Nevada Rosy-Finch by 41-60% since 1954-1980, with further losses expected.

Consistent with previous research, we found that the Brown-capped Rosy-Finch, currently recognized as a separate species, constitutes a distinct ESU from ecological and genetic perspectives. Overall, our results suggest that this lineage is sister to the clade that includes the Sierra Nevada Rosy-Finch, but, unlike the Sierra Nevada Rosy-Finch, it is connected via gene flow to all other lineages in the complex (Figure 1e). Our habitat suitability and GEA analyses suggest that, in contrast to the Sierra Nevada Rosy-Finch, the Brown-capped Rosy-Finch occupies and is adapted to the interior Rocky Mountain climate, characterized by high elevation and high summer precipitation (Figure 1f & 4c). Increased sampling across the Brown-capped range also revealed a separate lineage nested within the Brown-capped group in the Pikes Peak region of Colorado. Despite its apparent distinctiveness in the phylogeny, our admixture analyses and previous landscape-genomic work by DeSaix et al. (2022) suggest that Pikes Peak represents one end of a continuum in gene-environment relationships rather than a discrete, ecologically distinct ESU. Overall, our results help clarify how gene flow and local adaptation have shaped the genetic distinctiveness of the Brown-capped Rosy-Finch.

Although including samples from every major species and subspecies within this complex helped resolve relationships, additional work is needed, especially in the northern regions. Consistent with previous studies, our analyses suggest that the Black Rosy-Finch (*L. atrata*), currently recognized as a distinct species (Johnson et al., 2020), forms a monophyletic group. Notably, by including samples from the Wallowa subspecies (*L. t. wallowa*) of the Gray-Crowned Rosy-Finch, we find that it does not cluster with other Gray-Crowned Rosy-Finches as expected, but instead is sister to the Black Rosy-Finch (Figure 1a). Although this result contrasts with the current taxonomy, it is consistent with previous research attributing the darker plumage of Wallowa birds to historical admixture with the Black Rosy-Finch (Miller, 1939). In contrast to Funk et al. (2021), who found that the insular forms (*L. t. umbrina & L. t. griseonucha*) were strongly divergent from the rest of the Gray-crowned Rosy-Finch, we find that they are a distinct lineage nested within the Gray-crowned Rosy-Finch clade. We also observed little difference between the coastal Hepburn’s (*L. t. littoralis*) and the interior (*L. t. tephrocotis*) Rosy-Finch subspecies, although this may reflect our more limited sampling. Future work should prioritize expanded sampling across the northern parts of the range to clarify patterns of divergence, explore the uniqueness of the insular forms, and assess the extent to which the Wallowa lineage warrants recognition as a separate ESU.

### Evolutionary Potential

While adaptive capacity is a crucial part of climate vulnerability assessments, it is often overlooked or poorly defined (Thompson et al., 2015; Thurman et al., 2020). In general, adaptive capacity is an organism’s ability to respond to environmental change through dispersal, phenotypic plasticity, or genetic change; the latter is also referred to as evolutionary potential (Foden et al., 2019; Forester et al., 2022). A comparison between the Sierra Nevada and Brown-capped Rosy-Finch provides a useful context for assessing how adaptive capacity varies across alpine systems, as the two ESUs differ in several important ways. First, our analyses support the idea that the Sierra Nevada Rosy-Finch occupies a smaller area of suitable habitat (3,739 km2 in SNRF versus 13,929 km2 in BCRF), exhibits greater environmental heterogeneity (95% PCA ellipse area in SNRF was 37.22 vs. 18.65 in BCRF), and has more limited population connectivity than the Brown-capped Rosy-Finch (Figure 2e, 4c & 4d). Second, while both are altitudinal migrants (Grabherr et al., 2010; Lou & Therkildsen, 2022), only the Brown-capped Rosy-Finch overlaps with all other lineages, except the Sierra Nevada Rosy-Finch, during winter, thereby providing greater potential for dispersal. In addition, of the two southern lineages, the Brown-capped Rosy-Finch is geographically closer to other North American Rosy-Finch taxa during breeding summers, potentially further facilitating gene flow. We investigate how these key differences influence their adaptive capacity and ultimately contribute to differences in climate vulnerability.

Standing genetic variation can serve as a reliable indicator of evolutionary potential and, in some cases, may be a more accurate predictor of the capacity to evolve in response to climate change than adaptive genetic variation (Fournier-Level et al., 2016; Kardos et al., 2021). We found a northward decrease in genetic diversity (measured as H_IND_) from Brown-capped Rosy-Finches in the south to Gray-crowned Rosy-Finches in the north, except for the Sierra Nevada Rosy-Finch, which exhibited the lowest heterozygosity across the complex, despite its southerly distribution (Figure 4a). A decrease in heterozygosity with increasing latitude may reflect a decrease in genetic diversity with increasing distance from the center of a population expansion (Slatkin & Excoffier, 2012), and/or increased gene flow between Brown-capped Rosy-Finches and their northern relatives during the breeding and non-breeding periods, as discussed above (Figure 1e & 2a). In contrast, low heterozygosity in Sierra Nevada Rosy-Finches may result from their relative geographic isolation in the Southwestern part of the range, compounded by a recent decline in suitable habitat and a corresponding reduction in effective population size over the last 100 years (Figure 3c; Amirkhiz et al., 2025). Interestingly, heterozygosity has remained relatively high in the Brown-capped Rosy Finches, despite a similar recent decrease in suitable habitat and effective population size (Figure 3c; Amirkhiz et al., 2026). Overall, lower heterozygosity across Sierra Nevada Rosy-Finch populations indicates that it possesses lower evolutionary potential than Brown-Capped Rosy-Finches, which breed at similar latitudes.

Landscape genomic methods offer an additional way to evaluate evolutionary potential within populations by identifying genetic regions closely associated with environmental variation and thus potentially under selection (Rellstab et al., 2015). Compared with standard genome-wide genetic diversity metrics, these methods enable researchers to pinpoint adaptive genetic variation that may be important for a population’s capacity to evolve in response to climate change (but see Kardos et al. 2021). Although adaptive genetic variation often correlates positively with habitat heterogeneity, the ability of populations to adapt to local conditions depends, in part, on the amount of standing genetic variation (Forester et al., 2022). We demonstrated that, despite significantly higher habitat heterogeneity (17% greater; Figure 4c), the Sierra Nevada Rosy-Finch exhibits lower standing genetic variation at all loci, including those linked to local adaptation (Figures 4a & 4b). In fact, the Sierra Nevada Rosy-Finch shows the lowest levels of heterozygosity at putatively adaptive loci, which may result from past environmental selection driving allele frequencies to fixation but may also hinder its ability to adapt to future climate shifts. While previous findings by Robertson et al. (2026) support the idea that Sierra Nevada Rosy-Finch populations from sites with contrasting elevational profiles are distinct at loci associated with traits thought to be important to local adaptation (bill morphology and feather microstructure), the capacity for further evolution in these traits may be limited by this lineage’s overall low genetic diversity at both putatively adaptive and neutral loci.

Potential limitations in the Sierra Nevada Rosy-Finch’s ability to respond to future climate change are further highlighted by our genomic offset models under a moderate climate-change emissions scenario (SSP2-45). Genomic offset across the complex was predicted to be very low, but this likely reflects an inaccurate range map, which overestimates offset in regions along the Alaskan Coast and skews the overall picture. In contrast, when we employ high-resolution spatial maps for the Brown-Capped and the Sierra Nevada Rosy-Finch, it becomes clear that the Sierra Nevada Rosy-Finch will likely encounter more pressure to shift allele frequencies to keep pace with climate change than the Brown-Capped Rosy-Finch (Figure 4b). Higher genomic offset in the Sierra Nevada Rosy-Finch may be partly due to their observed lower evolutionary potential, combined with greater expected changes under climate change in the drier, more Mediterranean climate to which they are adapted.

### Effective & Census Population Size

While effective population size helps clarify how genetic drift and inbreeding influence long-term evolutionary potential, census population size helps clarify how ecological factors, such as range size and habitat availability, influence short-term demographic resilience. We estimate the effective population sizes of the Brown-capped and Sierra Nevada Rosy-Finch using a linkage-disequilibrium-based genetic method and compare them with recent estimates of census population size obtained via range-wide distance sampling (Bernier et al., 2023; Brown et al., 2026). Based on our analysis of effective population size over time, both lineages show evidence of recent declines (Figure 4c), although the timing of these declines should be interpreted cautiously given uncertainty in generation time. While the current effective population size of both lineages falls above commonly cited minimum thresholds of 500 to 1000 individuals needed to maintain evolutionary potential (Frankham et al., 2014; Franklin, 1980; Franklin et al., 2014; Lande & Barrowclough, 1987), the effective population size of the Sierra Nevada Rosy-Finch was estimated to be larger (5,841, 95% quantile: 4230 - 8,063) than that of the Brown-capped Rosy-Finch (2,364, 95% quantile: 1,711 - 2,952), even when we add the Pike’s Peak population to the total (78 + 2,364 = 2,442). Notably, this pattern was reversed for census population size, where the census size of the Sierra Nevada Rosy-Finch (32,700 to 50,800 individuals; Brown et al., 2026) was estimated to be smaller than that of the Brown-capped Rosy-Finch (67,600 to 149,000 individuals; Bernier et al., 2023), as expected based upon its larger breeding range (SNRF = 3739 km^2^ and BCRF = 13,929 km^2^; Figure 3d).

There are a couple of reasons the effective population size of the Brown-capped Rosy-Finch may be smaller than expected, given its range size. First, discrepancies between genetic and census estimates of population size may result from known variation in sex ratios among lineages (Bailey, 1974; French, 1959; Johnson, 1965; King & Wales, 1964; Shreeve, 1980; Twining, 1938), which, if they are heavily skewed, could reduce the effective size of the Brown-capped Rosy-Finch relative to its census size. Second, the linkage-based method we employed is known to yield downwardly biased estimates of effective population size in the presence of admixture (Santiago et al., 2020), which our results suggest is ongoing between the Brown-Capped Rosy-Finch and most other lineages in the complex (Figure 2). Together, these results suggest that although both ESUs currently maintain effective population sizes above commonly cited thresholds for sustaining evolutionary potential, past and projected changes in population size indicate they remain vulnerable to future declines in suitable habitat.

### Adaptive Capacity + Sensitivity + Exposure = Climate Vulnerability

Having quantified adaptive capacity for the Sierra Nevada and Brown-capped Rosy-Finches, we are now well-positioned to integrate information on exposure and sensitivity to provide a more complete assessment of climate vulnerability in the two southernmost lineages. Sensitivity is defined here as the degree to which a species is affected by or susceptible to climate change, is based on factors such as habitat specialization, physiological tolerances, and interspecific dependencies (IPCC, 2014). In turn, exposure is defined as the magnitude of climate change a species has experienced or is projected to experience (IPCC, 2014). While the two are linked, we begin by describing evidence for each lineage’s sensitivity to climate change, then discuss how this sensitivity may be amplified by differences in exposure and adaptive capacity.

While strong reliance on snowfields for summer foraging (Bernier et al., 2023; Brown, 2026; Brown et al., 2026; Epanchin, 2009) leads to the prediction that both lineages will be highly sensitive to climate-driven changes in snowpack, this sensitivity is further supported by both our landscape genomic analyses and previous ecological modeling efforts of Amirkhiz et al. (2025, 2026). Specifically, our landscape genomic analyses suggest that genetic variation in Brown-Capped and Sierra Nevada Rosy-Finches is most strongly associated with differences in soil moisture, mean summer precipitation, relative humidity, and elevation, all of which are linked to snowpack (Figures 1f and 4a). If these genetic-climate associations are important for fitness, we predict that changes in snowpack would lead to range shifts and/or population declines, particularly when range shifts are not possible. Correspondingly, recent ecological modeling work suggests that both species have shifted away from their former breeding habitats in the last 50-100 years, partly in response to changes in snowpack (Amirkhiz et al., 2025, 2026). In addition, our genetic estimates of effective population size over time suggest that both lineages have experienced population declines over roughly the same period as the projected changes in the availability of suitable habitat (Figure 3c), supporting the idea that snowpack loss may have fitness impacts. In summary, both our landscape genomic analyses and the ecological modeling work of Amirkhiz et al. (2025, 2026) support the conclusion that Sierra Nevada and Brown-capped Rosy-Finches are and will continue to be highly sensitive to climate change.

Alpine species generally face increased exposure to climate change because higher elevations are warming more rapidly than lower elevations (Mountain Research Initiative EDW Working Group, 2015). Although both species are losing snowpack, the Sierra Nevada—with its milder, marine-influenced winter temperatures—has lost, and is predicted to continue to lose, summer snowpack more rapidly than the Rocky Mountains (Siirila-Woodburn et al., 2021), suggesting it may have higher exposure levels. Specifically, April 1 snow-water equivalent, the primary proxy for June-August Rosy-Finch breeding-season snowpack, has declined by 5-40% at most Colorado Rocky Mountain and Sierra Nevada sites since 1955, with many Colorado sites experiencing up to 60% loss (USGCRP, 2023). By 2030-2100, April 1 snow-water equivalent is projected to decline by approximately 40-80% in the Sierra Nevada and 20-60% in the Rocky Mountains (Beltran-Peña et al., 2025; Siirila-Woodburn et al., 2021). Further, a recent model predicts that exceptionally low-snow years are becoming more severe (Belmecheri et al., 2016), with persistent low-to no-snow conditions in California from 2057 and in the Upper Colorado River Basin from 2079 (Siirila-Woodburn et al., 2021). This higher exposure to climate change in the Sierra is further supported by our genomic offset analyses, which predict that the Sierra Nevada Rosy-Finch will face greater pressure to undergo genetic changes to keep pace with climate change than the Brown-capped Rosy-Finch (Figure 4b). In summary, although both lineages are highly sensitive to climate-driven snowpack losses, the Sierra Nevada Rosy-Finch has lower adaptive capacity and will continue to face greater exposure to future climate change.

### Conservation Recommendations

At present, the most significant threat to Rosy-Finches on their summer breeding grounds is climate change and its associated effects, including extreme weather. Therefore, conservation of the Sierra Nevada and Brown-Capped Rosy-Finches will need to include climate change mitigation. While climate mitigation can lessen, it cannot fully prevent ongoing declines in their distribution and numbers due to climate system inertia. Because immediate, large-scale climate change mitigation is unlikely and beyond the scope of conservation efforts, effective strategies include enhancing their adaptive capacity through evolutionary interventions such as genetic rescue and ecological actions that increase resource availability for breeding Rosy-Finches. To increase the evolutionary potential of the Sierra Nevada Rosy-Finch, options might include genetic rescue from populations that more closely match its anticipated future needed genetic makeup. In addition, in California, removing introduced fishes from and stopping trout stocking in historically fishless alpine lakes could reduce the Sierra Nevada Rosy-Finch’s reliance on summer snowpack for insects vital to rearing their young, especially in breeding basins with lakes (Clapp et al., 2025; Epanchin et al., 2010). While resource enhancement options are limited in Colorado, restoring fishless lakes could also benefit Brown-capped Rosy-Finch populations breeding in drier, lower-snowpack regions.

## Conclusions

In conclusion, we demonstrate how integrating ecological and evolutionary data can yield a clearer understanding of the climate vulnerability and conservation needs of alpine species. Through increased sampling and genomic analysis, we provide the clearest picture to date of the evolutionary relationships within this iconic group, emphasizing the Sierra Nevada Rosy-Finch as an Evolutionary Significant Unit that is highly vulnerable to future climate change and warrants protection. More generally, this work demonstrates that combining ecological and genomic data can improve assessments of adaptive capacity across alpine systems. In turn, better estimates of adaptive capacity help clarify how it interacts with the other two components of climate vulnerability—sensitivity and exposure—to guide conservation priorities under climate change.

## Acknowledgments

We thank Sarah Albright, Alena Arnold, Emma Brown, Carl Brown, Alison Brunschwiler, Vincent Chevreuil, Ryan Cheung, Daisy Cortes, Jessica Gallardo, Karim Hanna, Raquel Lozano, Juan Rebellon, Leticia Santillana, Kevin Silberberg, and Juyung Yoo for their participation in catching, handling, banding, measuring, and sampling birds. Funding for this project was provided by the National Science Foundation (NSF 2222524, NSF 2222525, and NSF 2222526) and the American Ornithological Society. We thank NovoGene for their assistance with Next-Generation sequencing and Christine Rayne and Amanda Carpenter for library preparation. This work utilized the RMACC Summit supercomputer, which is supported by the National Science Foundation (awards ACI-1532235 and ACI-1532236), the University of Colorado Boulder, and Colorado State University. We also thank the University of Washington Burke Museum (UWBM), the Museum of Southwestern Biology (MSB), the Museum of Vertebrate Zoology (MVZ), the Yale Peabody Museum (YPM), the University of Alaska Museum (UAM), the KU Biodiversity Institute and Natural History Museum (KU), and the Denver Museum of Nature and Science (DMNS) for providing tissue and blood samples. Figures were created with https://BioRender.com, and illustrations were created by Erica Robertson. Handling, banding, and sampling of the Sierra Nevada Grey-crowned Rosy Finch were performed under Federal Banding Permit 24401 and CFWScientific 688 Collecting Permit (SCP) S-190610001-22081-001-02.

## Author Contributions

E.Z., K.R., and M.H. conceived the study and provided funding and resources. K.R., E.Z., and C.B. co-wrote the manuscript. C.B. created the figures with help from N.G. and R.G. T.B., E.Z., K.B., and E.R. performed fieldwork. C.B. analyzed the samples and genetic data. B.V., P.E.B., E.F., S.A.T., contributed samples and genetic expertise. All authors edited and approved the final manuscript.

